# Complete Chloroplast Genome of a Montane Plant *Spathoglottis aurea* Lindl.: Comparative Analyses and Phylogenetic Relationships among members of Tribe Collabieae

**DOI:** 10.1101/2023.09.10.557090

**Authors:** Nurul Shakina Mohd Talkah, Jasim Haider Mahmod Jasim, Farah Alia Nordin, Ahmad Sofiman Othman

## Abstract

The yellow–flowered *Spathoglottis aurea* (tribe Collabieae; family Orchidaceae) is native to the mountainous areas of Peninsular Malaysia. The species is well known as an ornamental plant and for its role in artificial hybrid breeding. There is an interesting evolutionary relationship between *S. aurea* and the geographically isolated *S. microchilina* from Borneo that has encouraged further study of the *S. aurea* populations, but the genomic resource for *S. aurea* has not yet been reported. The present study reports the first work to characterize a chloroplast genome among the *Spathoglottis* genus. The complete chloroplast (cp) genome of *S. aurea* was assembled from a sequence generated by the Illumina platform and analysed in comparison with other Collabieae species available in the GenBank database. The cp genome of *S. aurea* is 157,957 base pairs (bp) in length with guanine-cytosine (GC) content of 37.3%. The genome possessed a typical quadripartite chloroplast genome structure with large single-copy (LSC) (86,888 bp), small single-copy (SSC) (18,125 bp) and inverted repeat (IR) (26,472 bp) sequences. A total of 134 genes were annotated, with 88 protein coding genes (PCGs), 38 transfer RNA (tRNA) genes and eight ribosomal RNA (rRNA) genes. Overall, 80 simple sequence repeats (SSR) or microsatellites were identified. Comparative analysis with other Collabieae species revealed high conservation in the cp genome arrangements with minimal difference in genome lengths. However, several mutational hotspots were also detected, with high potential to be developed as genetic markers for phylogenetic analysis. Characterization of the *S. aurea* cp genome revealed its conserved nature without gene loss or rearrangements when compared to other species of the Collabieae tribe. Phylogenetic analysis of Collabieae species also revealed that *S. aurea* has a distant evolutionary relationship to other members of the Collabieae species, despite the presence of problematic genera such as *Phaius* and *Cephalantheropsis*.

## Introduction

The chloroplast genome is about 120-200 kilobases (kb) in size. In term of gene size, gene content, gene arrangements and genomic structure, it has frequently been shown to be conserved, except for a number of taxa (1,2). These characteristics, together with the absence of recombination and inheritance through a single lineage, make it a suitable marker for phylogeny and genetic variability studies of terrestrial plants. It has been shown to be a powerful molecular tool for taxonomy study (3), phylogenetics (4,5) and conservation genetics.

Collabieae is a tribe of orchids within the large family of Orchidaceae. It was first described by Pfitzer in 1887 (6), based on the genus *Collabium* and predominantly consists of terrestrial orchids. The high diversity of morphological characteristics such as vegetative and floral variations within Collabieae has made it difficult for systematists to infer their evolutionary relationships. Despite this, based on a revision by Chase (7), among the Collabieae tribe the chloroplast genome has been reported for several taxa, such as *Calanthe* (8–11) and *Tainia* (12–14). Although the group has approximately 450-500 known species, scarce genomic data are available and there are few reports of taxonomic conflicts (9,15,16).

The genus *Spathoglottis* Blume of Collabieae tribe is a favourite of horticulture enthusiasts, because it has bright, colourful flowers in deep crimson, pink and golden yellow that bloom all year round, is easy to cultivate and quick to grow. The Plants of the World Online (POWO) database, facilitated by Kew Gardens, contains 49 accepted species of the *Spathoglottis* genus. The natural distribution of this genus has been recorded in the Pacific Islands and in tropical and subtropical Asia. Across the Malesian region there are 44 species that are widely distributed, with particularly dense gatherings in New Guinea (15). There are four *Spathoglottis* species recorded in Peninsular Malaysia with one infraspecific taxon: *S. aurea* Lindl., *S. plicata* Blume, *S. hardigiana* C.S.P.Parish & Rchb.f., *S. affinis* de Vriese and *S. plicata* var. *alba*. The first of these is a common mountain species native to Peninsular Malaysia, whose population is restricted to the top of mountains, where it usually grows on the granitic terrain of moist forests. It was first described by Lindley in 1850, based on a specimen from Mount Ophir (Gunung Ledang) in Johor (17) and was named *S. wrayi* 40 years later by Hooker, before receiving its present-day name of *S. aurea*. As a part of the *Spathoglottis* genus with a yellow flower complex, *S. aurea* is unique because of the influence of anthocyanin in its floral parts and leaves. Recent molecular (15) and morphological (18) studies have shown that *S. aurea* could be the closest ancestor of *S. microchilina* Kraenzl., which has only been reported in the Borneo region. Further population studies may confirm this relationship between *S. aurea* and *S. microchilina*.

The present study aimed to: (a) characterize the cp genome of *S. aurea*; (b) compare the *S. aurea* chloroplast genome structure with that of fellow species of the Collabieae tribe; (c) determine the phylogenetic positions of *S. aurea* within the Collabieae tribe in terms of all chloroplast protein coding genes. The availability of the cp genome in our dataset is expected to support further study on *S. aurea* populations and the phylogeny of the *Spathoglottis* yellow flower complex.

## Materials and methods

### Sampling, DNA Extraction, and Sequencing

Individual *S. aurea* plants were collected from Mount Jerai, Kedah, Malaysia and replanted in a greenhouse at Universiti Sains Malaysia. The voucher specimen was deposited in the Herbarium, School of Biological Sciences, Universiti Sains Malaysia (USMP11942). Fresh leaves from the cultivar were cleaned with running tap water and 70% ethanol and put in silica gel to dry. Genomic DNA was extracted from the dried leaves using 2X cetyltrimethylammonium bromide (CTAB) according to the conventional method (19), with three modifications: (1) At the beginning of the protocol, after grinding the dry leaf tissue with liquid nitrogen, 750 μl CTAB-free buffer [200 mM Tris-HCl pH 8.0, EDTA 50 mM and NaCl 250 mM] and 5 μl β-mercaptoethanol were added. The samples were kept at -20°C for 10 minutes, before being subject to a centrifuge at maximum speed for 5 minutes and the supernatant discarded. These steps were repeated two or three times until a colourless or slightly yellow supernatant was obtained; (2) The chloroform – isoamyl alcohol (CIA) mixture (24:1) was added three times rather than twice; (3) After adding ice-cold isopropanol, the mixture was stored at -20°C for 1-2 hours. The samples were then centrifuged at 12,000 rpm for 10 minutes, the supernatant discarded, and the DNA pellet deposited carefully. A 600 μL wash buffer [2.5 M sodium acetate (NaAc); 76% ethanol, pH 5.0] was added to the DNA pellet to be washed and left at room temperature for 30 minutes, then centrifuged at 8,000 rpm for 1 minute and the supernatant discarded. The wash step was repeated twice, then the DNA pellet was allowed to dry completely at room temperature for 5 hours. Finally, the dried DNA was resuspended with 80-100μL of TE Buffer and incubated at 55°C for 30 minutes in a water bath. The genomic DNA was sequenced for 150bp paired end reads via the Illumina Novaseq 6000 using the TruSeq Nano DNA kit according to manufacturer protocol.

### Chloroplast Genome Assembly, Assembly Validations, and Gene Annotation

De novo assembly of *S. aurea* chloroplast genome was done using GetOrganelle version 1.7.5. (20). The assembled chloroplast genome was validated by using coverage depth analysis. This analysis was done by mapping back the assembled genome to raw reads using ‘mem’ function of Burrows-Wheel aligner (21) and sorted using SamTools (22). Coverage depth for each nucleotide position in the assembled genome were retrieved using SamTools. The finalized chloroplast genome was then annotated using GeSeq (23) with Chloe function invoked. The open reading frame positions were confirmed by comparing them with reference genomes of closely related species using Genbank Blast function (24). The annotation table generated by GB2Sequin was edited according to the correct reading frame before being deposited in the Genbank. Visualization of *S. aurea* chloroplast genome was done from Genbank flatfile using CPGView platform (25).

### Sequence Repeat Analysis and Codon Usage

Simple sequence repeats (SSRs) of the *S. aurea* cp genome were identified using MISA (26) with the parameter of 10 repeat units for mononucleotides, five repeat units for dinucleotides, four repeat units for trinucleotides and three repeat units for tetranucleotides, pentanucleotides and hexanucleotides. REPuter (27) was then used to detect forward, reverse, complement and palindrome repeats in the cp genome. Protein coding gene (PCG) sequences were extracted using Python scripts from the GenBank file. The PCG sequences were analysed for relative synonymous codon usage (RSCU) using the Bioinformatics web server (https://www.bioinformatics.org/sms2/codon_usage.html) (28) developed by Stothard (2000) and MEGA-X (29).

### Comparative Analysis of Chloroplast Genome

The gene content and rearrangement in the *S. aurea* cp genome were compared with eight other Collabieae species available in the GenBank using progressiveMauve in Mauve version 2.4.0 (30). The genes available near inverted repeat boundaries were visualized using IRscope (31). Whole chloroplast genomes of eight Collabieae species were aligned using MAFFT v7.511 (32). The sequence matrix was utilized to evaluate nucleotide diversity (Pi) using DNaSP6 (33) with sliding window analysis of 600bp window length and 200bp step size. The nucleotide diversity threshold was calculated by the sum of the average and double the variance of nucleotide diversity (Pi) (34).

### Phylogenetic Analysis

Phylogenetic analysis was performed using 71 PCGs extracted from 29 different Collabieae species and one outgroup species (*Dendrobium sinense*) available in the GenBank. All PCGs were concatenated using MEGA-X and aligned by MAFFT v7.511. The best-fit model for sequence evolution was assessed using ModelFinder (35). Finally, a maximum likelihood (ML) tree was built based on the TVM+F+R2 model using IQ-Tree version 1.6.12 (36), operated on the Linux platform with 1000 bootstrap replicates.

## Results

### Characterization of the Chloroplast Genome of *Spathoglottis aurea*

Paired-end raw read sequencing of *S. aurea* produced 95,319,748 reads (7GB) with a GC content of 37.09% and a Phred score (Q30) of 86.48%. The raw reads were deposited in the Sequence Read Archive (SRA) of GenBank (accession number SRX1938371). A total of 652,534 reads were assembled to complete the chloroplast genome of 157,957 bp in length (GC content: 37.3%; GenBank accession number: OQ411079). The resulted assembly shows an average of 1,600X genome coverage (Figure S1) with a typical quadripartite structure of large single copy (LSC), small single copy (SSC) and a pair of inverted repeats (IR). Overall, 134 genes with 88 protein coding genes (PCG), eight ribosomal RNA (rRNA) genes and 38 transfer RNA (tRNA) genes were annotated in. There are 18 genes with intron with two of the contain two introns respectively. The two genes contained two introns are *paf*I and *clp*P1. Details on other genes present in *S. aurea* cp genome and their functionalities are listed in Table 1. The visualization of circularized chloroplast genome is shown in Figure 1.

**Table 1.**
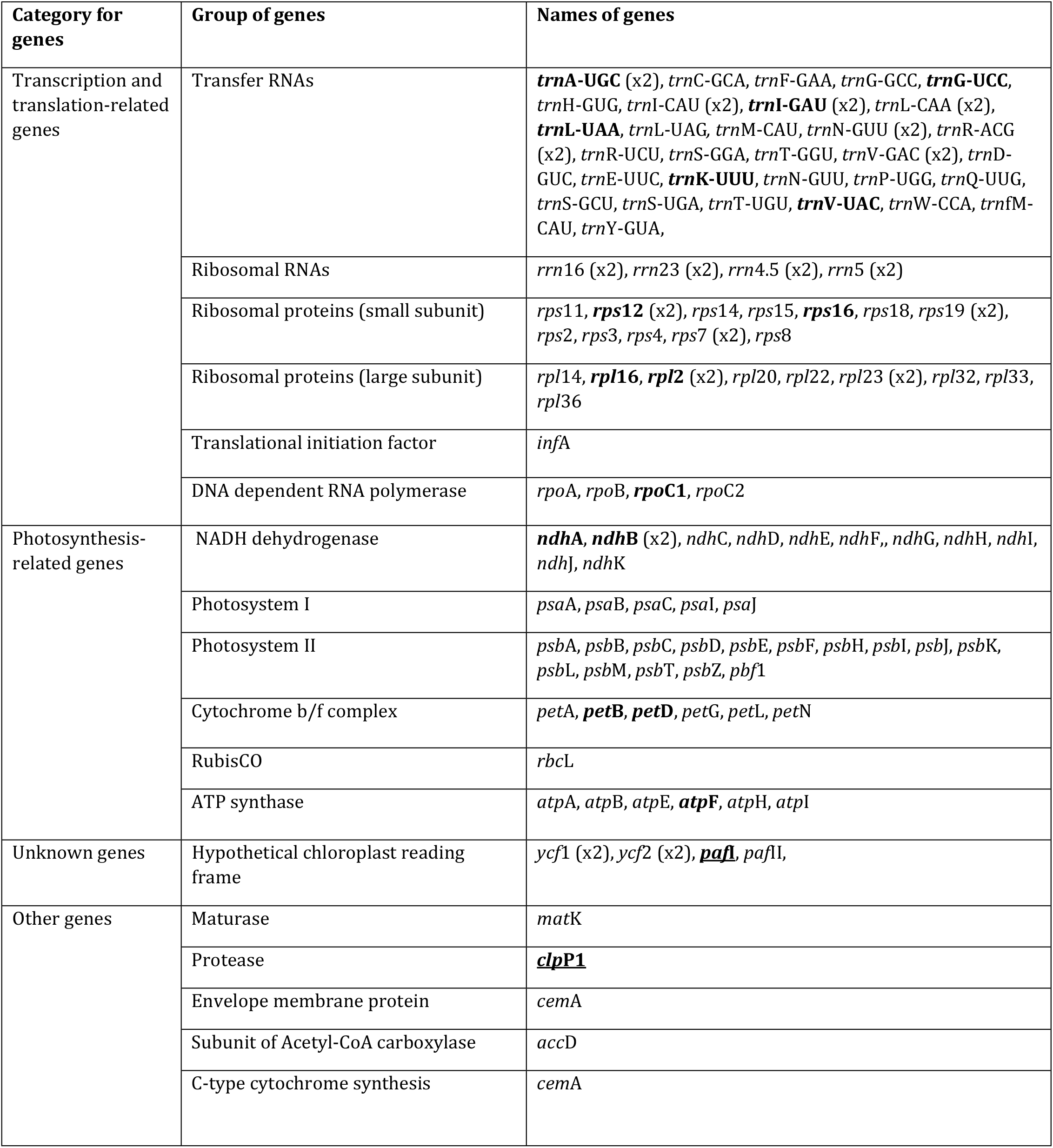
Genes present in the *S. aurea* chloroplast genome. Genes with bold letters: contain intron, genes with bold and underlined letters: contain two introns.

**Fig. 1.**
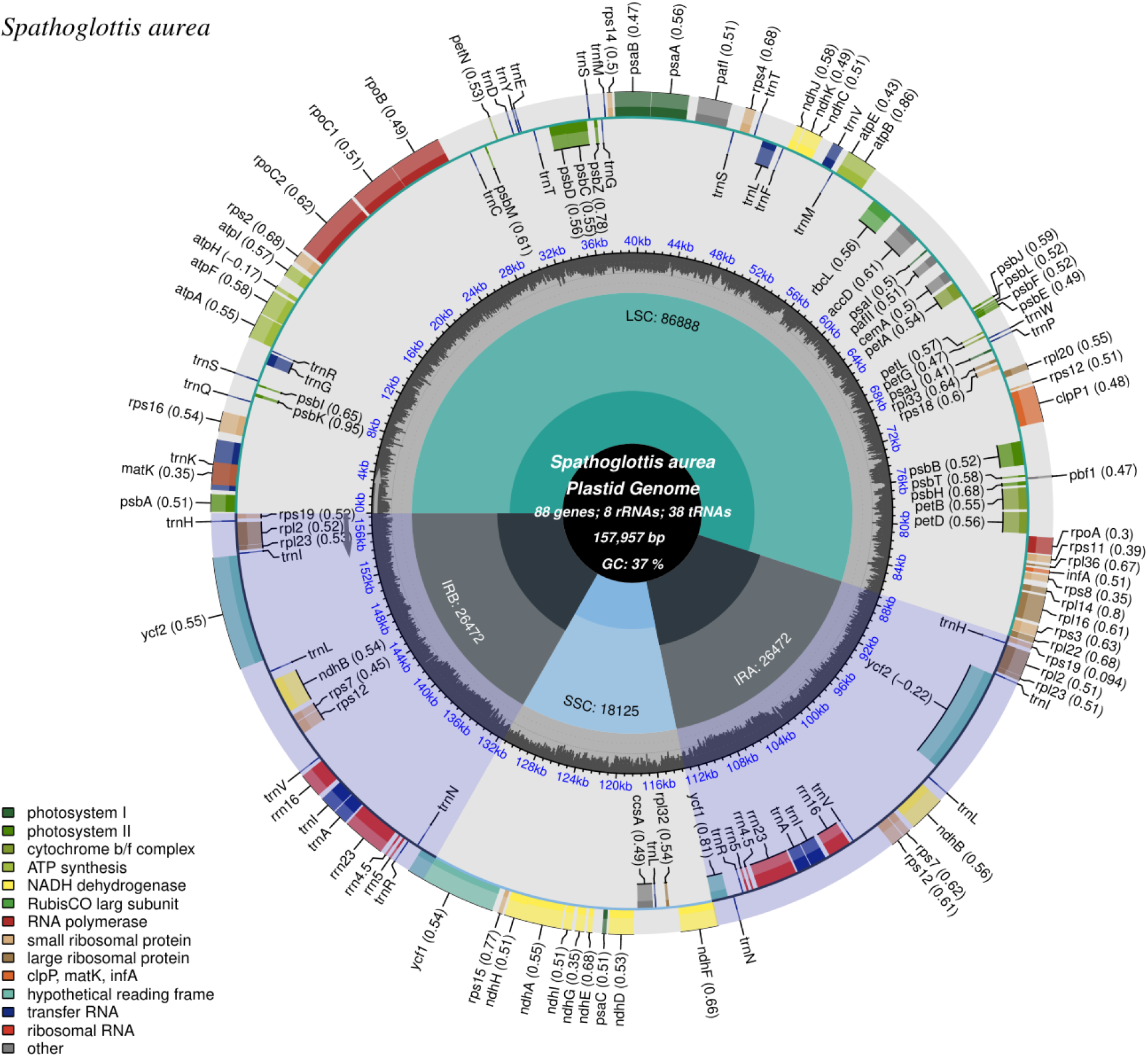
The figure shows visualization of *S. aurea* chloroplast genome (OQ411079) generated by CPGView. The small single-copy (SSC), inverted repeat (IRa and IRb), and large single-copy (LSC) regions were labelled respectively. The genes shown on the most outer track were color coded according to their functional groups with legends available on the left bottom. The genes that transcribed in clockwise direction positioned in inner lines while the genes that transcribed in anticlockwise directions are positioned in outer lines. The codon usage bias is displayed in the parenthesis after the gene name.

The *S. aurea* cp genome contains 13 protein coding cis-splicing genes, of which two have two introns (*paf*I; also named *ycf*3 and *clp*P1). The trans-splicing genes, *rps*12, consist of exons from three different regions; the 5′-end exon was found in the LSC region and the intron, and the 3′-end exon was located in the IR region. Figure 2 shows the list of cis-splicing and trans-splicing genes with their positions in the genome. There are 20,690 codons in the cp genome (Table S1). Codon usage analysis from *S. aurea* PCGs revealed codons ending with A/T nucleotides generally have higher usage compared to those ending with G/C nucleotides (Figure 3). The amino acid that was most coded in the genome is Leucine, with 2,118 codons. The codon occurring the most is GAA which is 871 times.

**Fig. 2.**
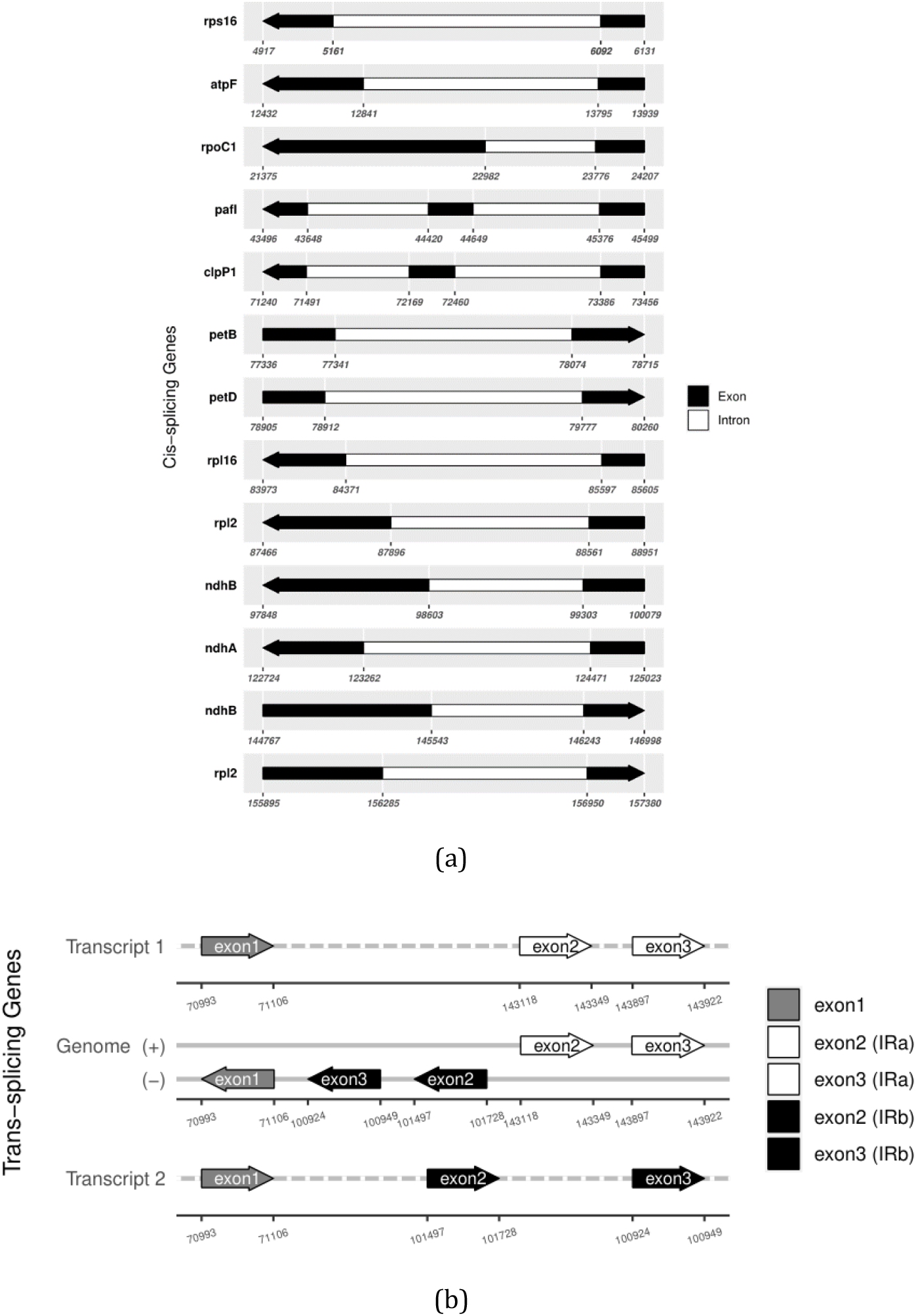
The figure shows genes splicing positions in *S. aurea* chloroplast genome. (a) The genes with cis-splicing regions were arranged according to their positions in chloroplast genome. (b) Trans-splicing gene *rps*12 with three unique exons. The gene is duplicated in inverted repeat regions.

**Fig. 3.**
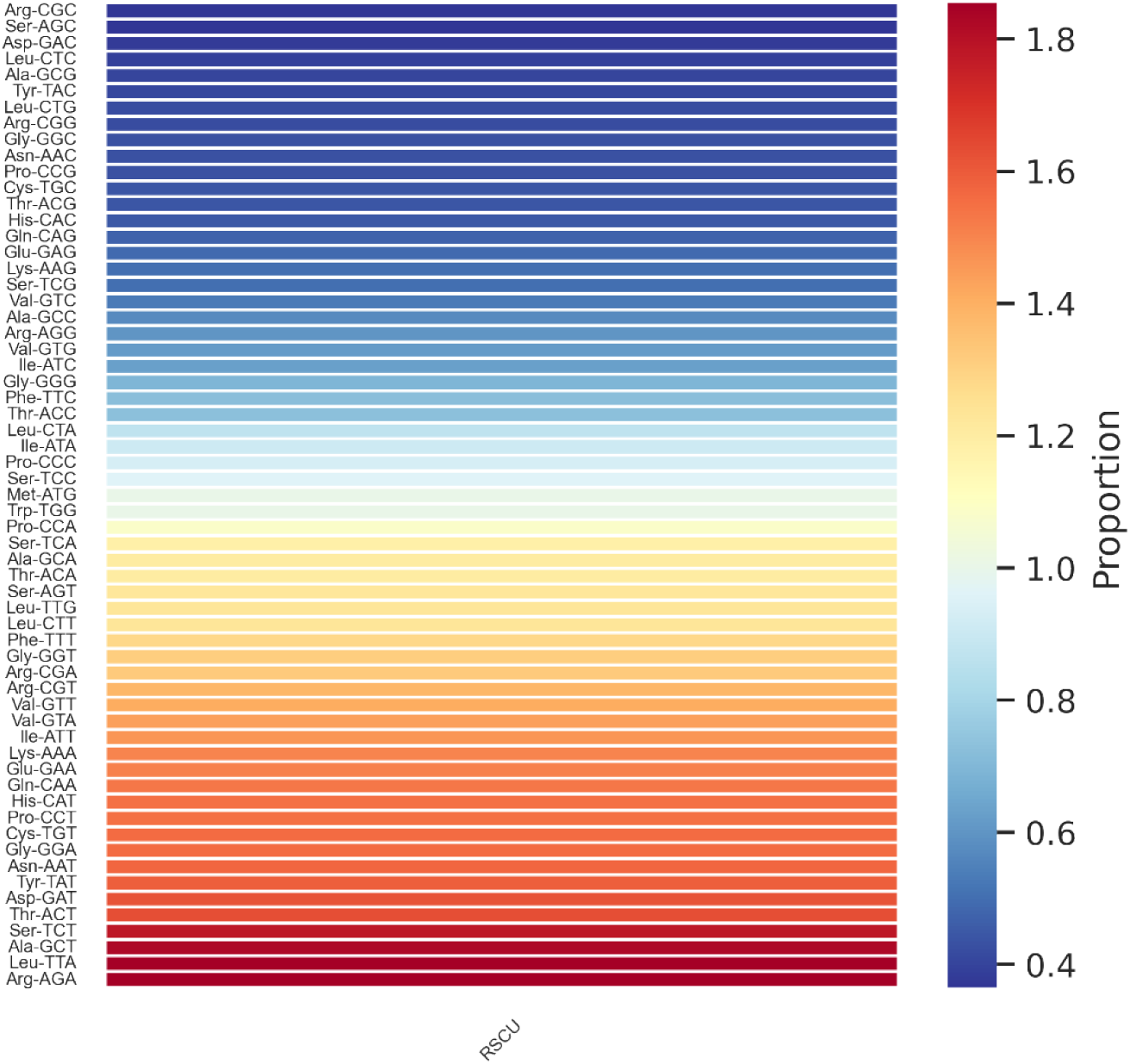
The figure depicts visualized relative synonymous codon usage in *S. aurea* chloroplast genome as heatmap.

**Fig. 4.**
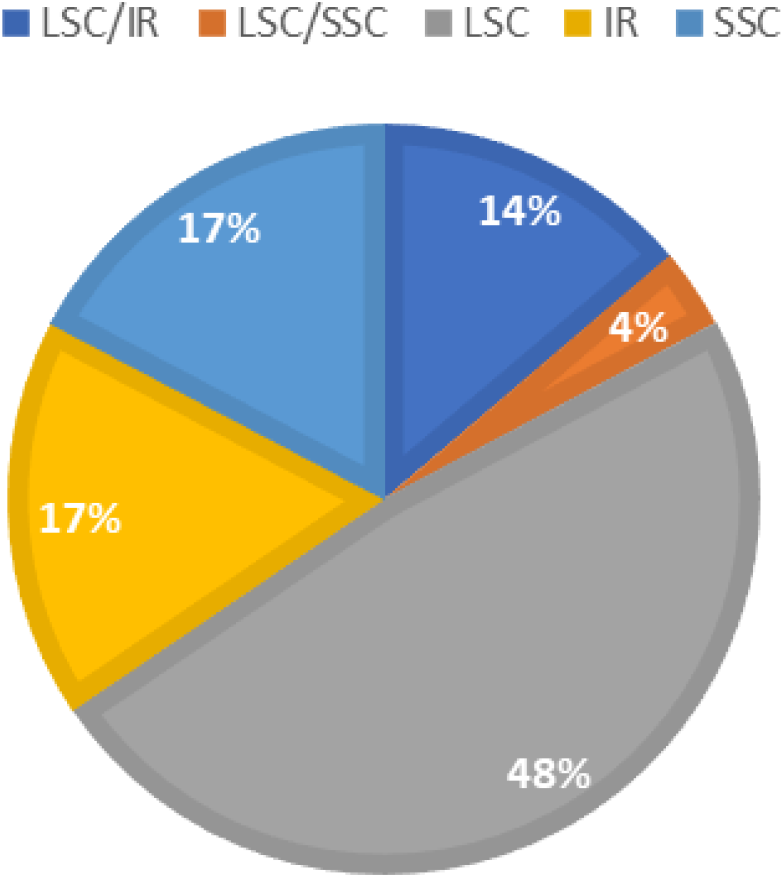
Percentage location of long repeats in the *S. aurea* cp genomes.

### Comparison of cp Genome Structures within Collabieae species

The lengths and gene contents for *S. aurea* and eight other Collabieae cp genomes are available in Table 2. Except for *Cephalantheropsis obcordata* and *Hancockia uniflora*, all other Collabieae species are widely distributed across Peninsular Malaysia. In general, all cp genomes show minimal variation in total length, ranging from 156,036 -158,828 bp. There is also minimal variation in length between LSC, SSC and IR. The total gene contents are conserved for all species.

**Table 2.**
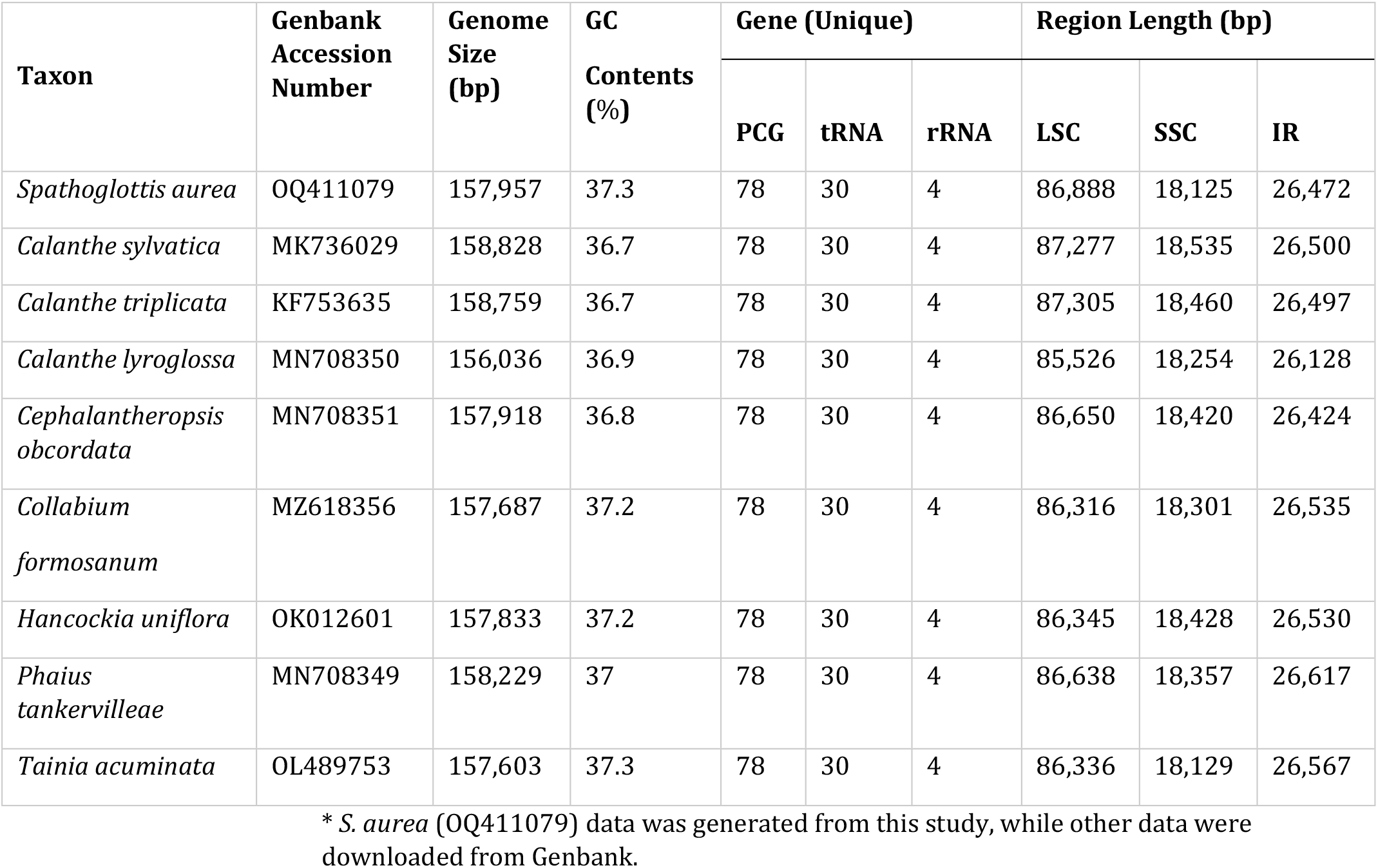
This table shows genomic features of 9 species in Collabieae tribe.

Gene rearrangements among Collabieae cp genomes are shown in Figure 5. The collinear block lines connecting all the cp genomes demonstrate similar gene arrangement, except for *Collabium formosanum*, in which the genes within the SSC region appear to be inverted.

**Fig. 5.**
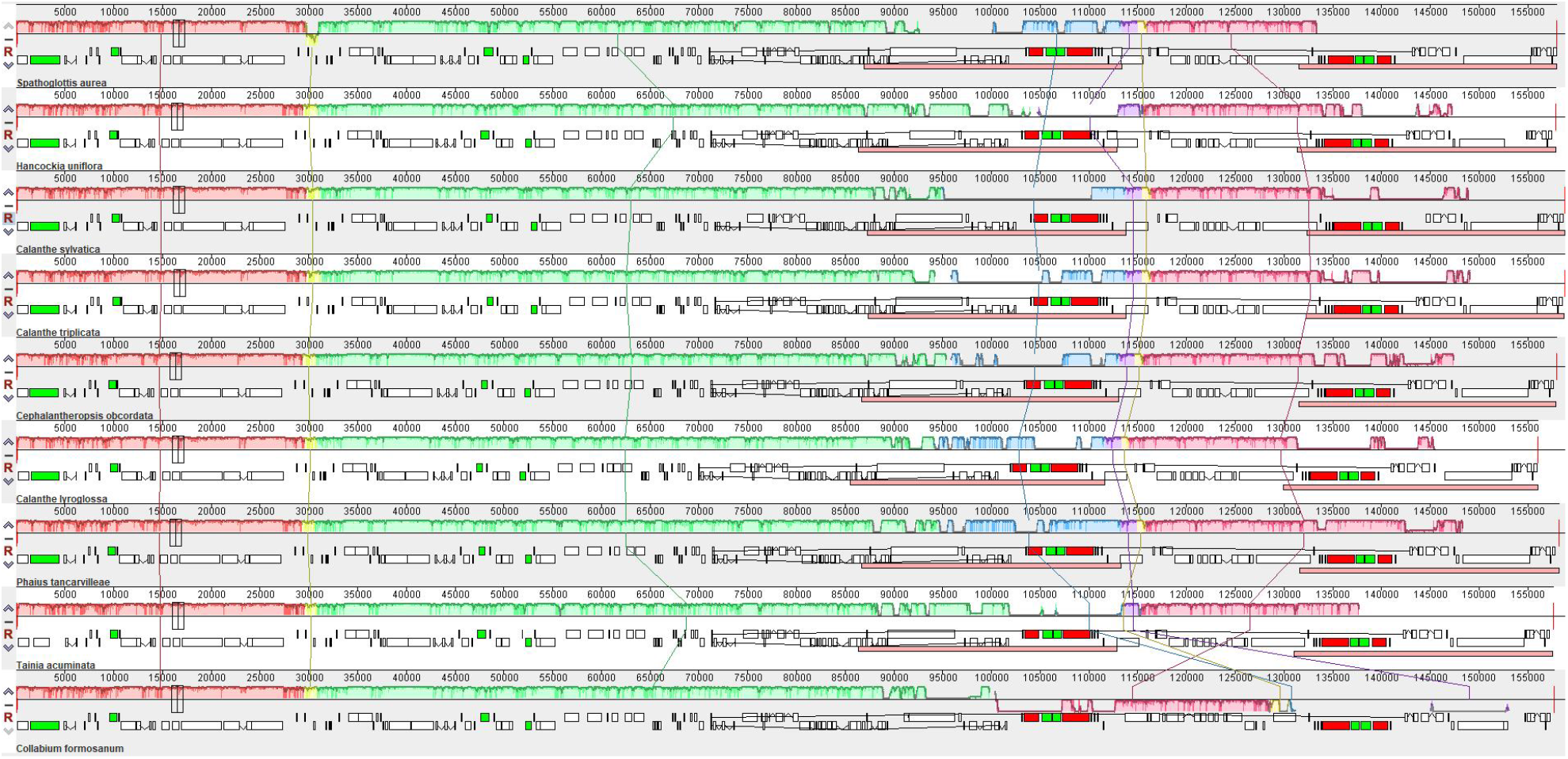
This figure was generated by progressiveMauve alignment which visualized any genes rearrangements among Collabieae tribe.

Within the eight Collabieae species, there are several variations at the junctions of LSC/IR/SSC. In *S. aurea* and *Calanthe lyroglossa*, the *rpl*22 gene is entirely located in the LSC region, at distances of 35 bp and 9 bp, respectively, from the JLB boundary. Conversely, for other species, *rpl*22 is located within the JLB boundary, with various lengths (Figure 6). The entire *ycf*1 gene is located within the JSB boundary for all species except for *Collabium formosanum*, which is located in the SSC region without overlapping any boundaries. Annotation from *S. aurea* and *Hancockia uniflora* resulted in truncated *ycf*1 to be observed in the IRa region near the JSA boundary. The *ycf*1 in these two species is part of a duplicated gene in which the complete gene copy was found at the JSB boundary. Likewise, the entire *ndh*F gene for *Collabium formosanum* is located within the SSC region, similar to that in *Hancockia uniflora*. For other species, *ndh*F is slightly expanded from the SSC to the IRa region, thus overlapping within the JSA boundary. For *rps*19, which is in both IR regions, the nucleotide distance from JLA boundaries varies from 59 – 463 bp.

**Fig. 6.**
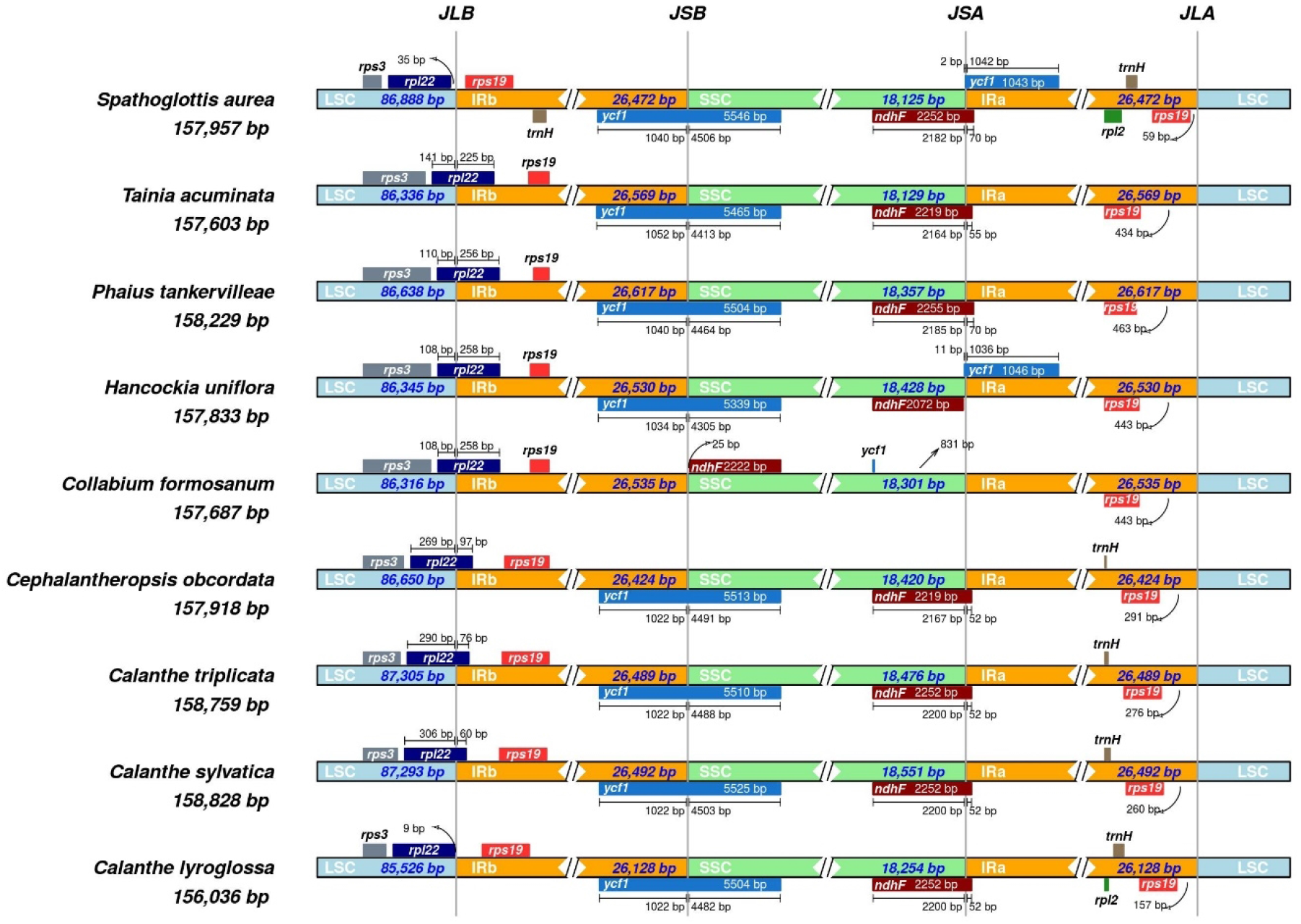
Comparison of LSC, SSC and IR junctions among 9 Collabieae species. Genes that are presented on top of the track are transcribed anti-clockwise while genes that are presented below of track are transcribed clockwise. The junctions are labelled as JLB (LSC/IRb), JSB (IRb/SSC), JSA (SSC/IgRa) and JLA (IRa/LSC).

For nucleotide diversity (Pi) analysis, *Collabium formosanum* was excluded from the data matrix because of its different SSC orientation. The threshold value for the data matrix is 0.04612; any regions with values higher than this are considered to have high nucleotide diversity. Seven regions demonstrated high nucleotide diversity, including *trn*K-UUU, *rps*16 – *trn*Q-UUG, *trn*C-GCA – *trn*Y-GUA, *ndh*C – *trn*V-UAC, *atp*B – *rbc*L and *psb*B-*psb*T (Figure 8). Such regions only occur in LSC and not in IR and SSC regions.

### Phylogenetic Analyses of cp within Collabieae

Further investigation of the phylogenetic position of *S. aurea* among Collabieae species was calculated based on the maximum likelihood (ML) algorithm (Figure 8). The outgroup for the tree is *Dendrobium sinense*, a species from the Malaxideae tribe. Clade A shows *S. aurea* [BP(ML) = 100%] bifurcate farther from other species. Clade B consists of *Collabium formosanum* with *Hancockia uniflora* as a sister clade [BP(ML) = 100%], which appeared to have a closer relationship to species in Clade C [BP(ML) = 100%] than other species involved in this study. All species of *Tainia* genus are clustered in Clade C. Finally, Clade D consists of the paraphyletic *Calanthe* genus along with two closely related genera, *Cephalantheropsis* and *Phaius*.

## Discussion

This study provides the first complete chloroplast genome for the *Spathoglottis* genus. The data will lead to the availability of omics repositories that can support further systematics, evolutionary, population and conservation research (15). Like most other angiosperm chloroplast, such as *Arabidopsis thaliana, S. aurea* possessed both cis- and trans-splicing genes. Cis-splicing genes are also referred to as normal gene splicing, whereby the joined exons originated from the same transcript. In contrast, trans-splicing genes require the joining of exons from different gene locations prior to mRNA processing (37). Other than the 13 PCGs in Figure 2, there are also six tRNA (*trn*G-UCC, *trn*L-UAA, *trn*V-UAC, *trn*I-GAU, *trn*A-UGC, *trn*V-GAC) genes and an rRNA (rrn23) gene that are cis-splicing genes in the *S. aurea* cp genome. *rps*12 is the most reported trans-splicing gene in the chloroplast genome and can exist in several configurations depending on the species (25). The configurations are based on the gene locations in the cp genome.

The development of microsatellites or SSR markers are very useful for populations and genetic relationship studies. Although SSR markers developed from the nuclear region give comprehensive information about both maternal and paternal parents, their high rate of sequence evolution makes them unsuitable for phylogenetic analysis of different taxa (38). SSR markers developed from the chloroplast region could be complementary here, because of this region’s highly conserved nature compared to that of the nuclear region (39).

Compared to other Collabieae species, the *S. aurea* cp genome shows a highly conserved structure without any gene loss. The quadripartite structure and gene compositions are consistent with previously reported Collabieae species (9,10,40). In Figure 5, gene inversion was observed in the *Collabium formosanum* cp genome because of its different orientation submitted in the GenBank (41). It is the nature of cp genomes to harbour two different structure orientations based on the SSC region. Thus, there is no gene rearrangement between Collabieae species in this study. Similarly, in Figure 6, the gene *ndh*F seems to be positioned differently from other species because of a different SSC orientation. Nevertheless, in *Collabium formosanum* there is no *ycf*1 gene available in the IR region because it is fully located inside the SSC region.

Nucleotide diversity (Pi) analysis of *S. aurea* revealed several mutational hotspots compared to seven other Collabieae species. Species from earlier analysis were used, except for *Collabium formosanum*. Different orientations of the SSC regions might lead to false interpretations. In Figure 7, seven regions are shown, which are mostly intergenic regions except for the *trn*C-GCA - *trn*Y-GUA region which also includes *psb*M, *pet*N and *trn*D-GUC as highly variable regions. These regions are useful for the development of molecular markers to resolve several taxonomic debates reported by previous studies (1,42,43).

**Fig. 7.**
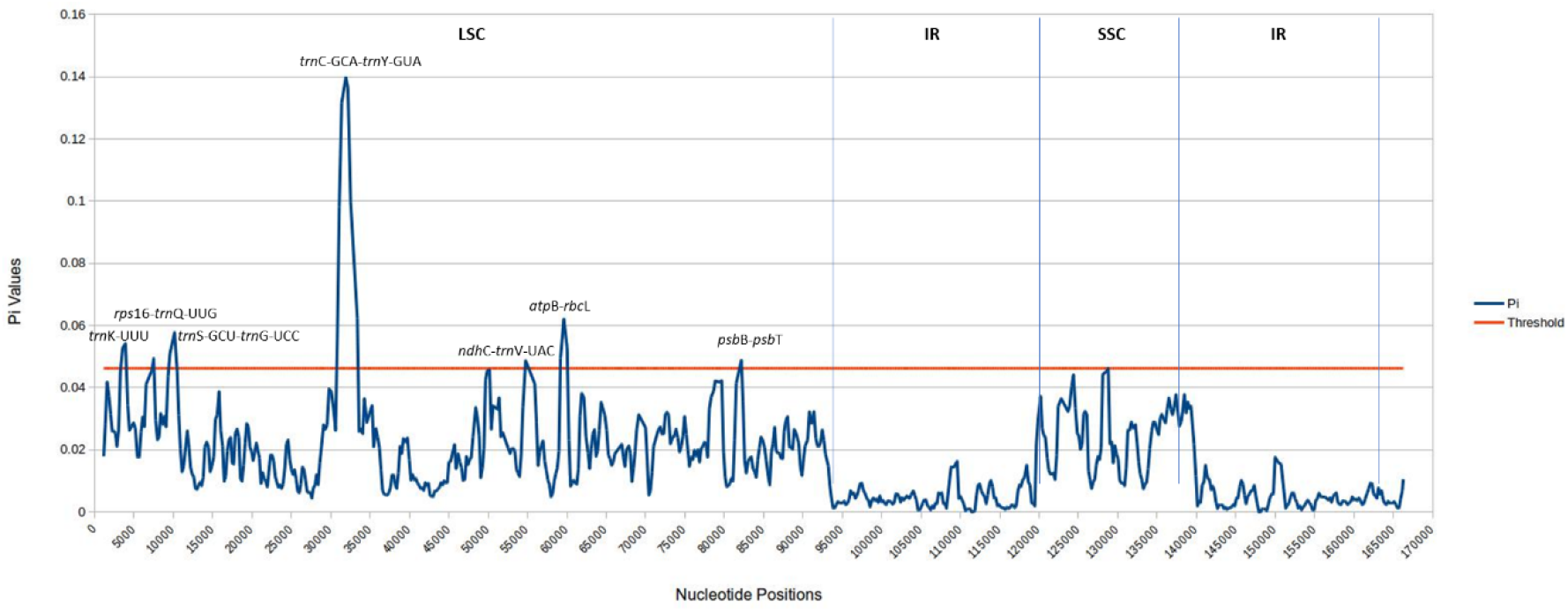
Nucleotide diversity (Pi) value of *S. aurea* chloroplast genome compared with several Collabieae species.

**Fig. 8.**
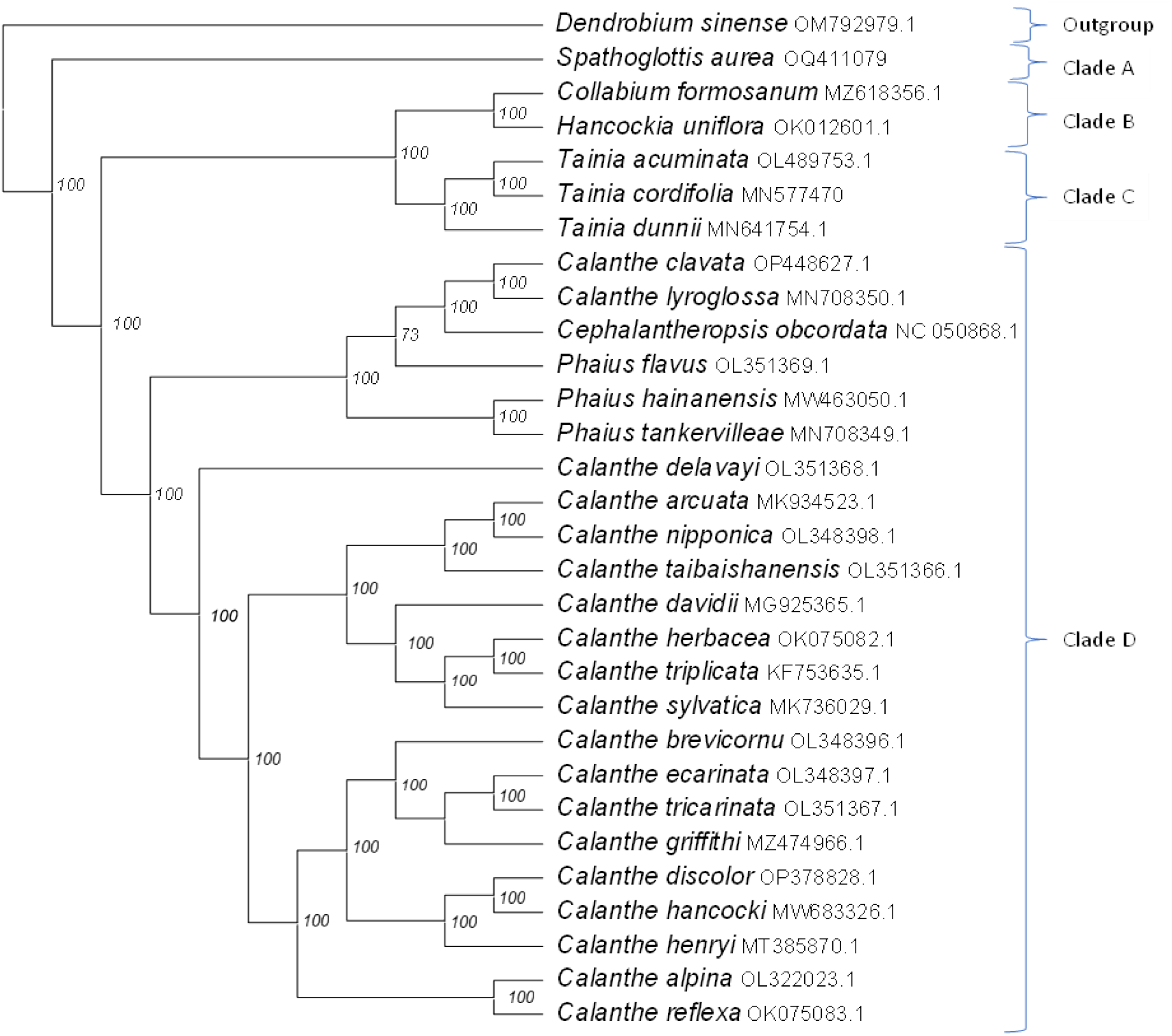
Phylogenetic tree was built by maximum likelihood (ML) algorithm using 71 PCGs shared by 29 Collabieae species and species outgroup (*Dendrobium sinense*).

The phylogenetic tree in Figure 8 revealed similar results to the molecular study by Xiang (44), which use four chloroplast genes (*mat*K, *psa*B, *rbc*L and *trn*H-*psb*A) to elucidate the relationship among Collabieae species. In the present study, *S. aurea* shows a further relationship with other species (Clade A); in particular, *Cephalantheropsis* and *Phaius* also reside in the same clade as *Calanthe*. Despite several differences in morphological characteristics, a number of molecular studies (9,44) including the present study have consistently shown that these two genera are positioned within the *Calanthe* clade. For the *Phaius* species, the morphological characteristic differences are minimal compared to those of the *Calanthe* species. However, species under the *Cephalantheropsis* genus showed significant differences in morphological characteristics compared to *Calanthe* and *Phaius*. Chase proposed to merge these two genera (*Cephalantheropsis* and *Phaius*) under *Calanthe* (16,45), but there is ongoing debate about this merger and it has not been universally accepted.

## Conclusion

This study characterized the complete chloroplast genome of *S. aurea*; one of the commonly known species used in landscaping and to develop artificial hybrids. The results provide the first genomic resource of the *Spathoglottis* genus. Data such as SSR locations will be useful for future assessment of *S. aurea* populations and other *Spathoglottis* species. The evolutionary relationship between *S. aurea* and other Collabieae species based on shared PCGs was also revealed to be consistent with previous studies, including the highly debated genera *Phaius* and *Cephalantheropsis* in Orchidacea.

## Supporting information

**S1 Fig. Coverage graph**. Coverage depth of nucleotide position of *S. aurea* chloroplast genome

**S1 Table. Codon table**. Codon usage table *S. aurea* chloroplast genome

**S2 Table. MISA file**. Original MISA file for SSR detection in *S. aurea* chloroplast genome

**S3 Table. SSR table**. Details on SSR location in the *S. aurea* chloroplast genome

## Acknowledgement

The study was funded by the USM Short Term Grant of assignment No. 304/PBI-OLOGI/6315549 awarded to the third author and the corresponding author. All funders provided financial supports for the present work but did not contribute any additional role in in the research design, data collections and analysis, and preparation of the manuscript.

## Author Contributions

**Conceptualization:** Ahmad Sofiman Othman, Nurul Shakina Mohd Talkah

**Data curation:** Nurul Shakina Mohd Talkah

**Formal analysis:** Nurul Shakina Mohd Talkah

**Funding acquisition:** Farah Alia Nordin, Ahmad Sofiman Othman

**Project administration:** Ahmad Sofiman Othman

**Resources:** Farah Alia Nordin, Jasim Haider Mahmod

**Software:** Nurul Shakina Mohd Talkah

**Writing - original draft:** Nurul Shakina Mohd Talkah, Jasim Haider Mahmod

**Writing - review & editing:** Ahmad Sofiman Othman, Farah Alia Nordin

